# Formin’s nucleation activity influences actin filament length

**DOI:** 10.1101/2021.06.01.446650

**Authors:** Mark E. Zweifel, Laura A. Sherer, Biswaprakash Mahanta, Naomi Courtemanche

## Abstract

Formins stimulate actin polymerization by promoting both filament nucleation and elongation. Because nucleation and elongation draw upon a common pool of actin monomers, the rate at which each reaction proceeds influences the other. This interdependent mechanism determines the number of filaments assembled over the course of a polymerization reaction, as well as their equilibrium lengths. In this study, we used kinetic modeling and *in vitro* polymerization reactions to dissect the contributions of filament nucleation and elongation to the process of formin-mediated actin assembly. We found that the rates of nucleation and elongation evolve over the course of a polymerization reaction. The period over which each process occurs is a key determinant of the total number of filaments that are assembled, as well as their average lengths at equilibrium. Inclusion of formin in polymerization reactions speeds filament nucleation, thus increasing the number and shortening the lengths of filaments that are assembled over the course of the reaction. Although variations in elongation rates produce modest changes in the equilibrium lengths of formin-bound filaments, nucleation constitutes the primary mode of monomer consumption over the course of assembly. Sustained elongation of small numbers of formin-bound filaments therefore requires inhibition of nucleation via monomer sequestration and a low concentration of activated formin. Our results underscore the mechanistic advantage for keeping formin’s nucleation efficiency relatively low in cells, where unregulated actin assembly would produce deleterious effects on cytoskeletal dynamics. Under these conditions, differences in the elongation rates mediated by formin isoforms are most likely to impact the kinetics of actin assembly.

## Introduction

Actin polymerization is a fundamental biological reaction that supports a broad range of essential cellular functions, including cell growth, division and motility. Actin assembly is tightly regulated in cells, both at the initial step of filament nucleation and during subsequent elongation. Spontaneous nucleation requires the assembly of energetically unstable actin trimers that become stable filaments through the association of an additional monomer [1–3]. A diverse cohort of actin-binding proteins control filament nucleation either through monomer sequestration or nucleation-promoting mechanisms, ensuring that filament assembly occurs at the appropriate time and subcellular location [4, 5]. Once assembled, filament nuclei continue to bind actin monomers at their barbed and pointed ends at rates that are modulated by elongation-promoting proteins [5].

Whereas many proteins that regulate actin assembly do so by influencing either filament nucleation or elongation, the formin family of proteins stimulates both processes [6–8]. Formins promote filament nucleation by encircling and binding actin nuclei via their dimeric Formin Homology 2 (FH2) domains [9–12]. This interaction stabilizes nascent actin nuclei and enables elongation through subsequent actin binding events [9, 11, 13–17]. Following nucleation, FH2 dimers remain bound at filament barbed ends and step processively onto incoming actin subunits to incorporate them into the filament [17]. Conformational fluctuations of the FH2 dimer “gate” the barbed end by regulating its availability for actin monomer binding, ultimately slowing elongation [17–19]. Formins overcome the effects of gating on elongation through transient interactions with the actin monomer-binding protein profilin [17, 18]. Profilin-actin complexes bind polyproline tracts located within formin FH1 domains [20, 21], enabling their rapid delivery to the barbed end via diffusion of these flexible domains [18, 22].

Most eukaryotes express at least two formin isoforms that assemble unbranched actin filaments that are incorporated into cytoskeletal structures including cytokinetic rings, filopodia and stress fibers [23, 24]. Each formin isoform mediates a specific rate of filament elongation that depends on both the extent to which its FH2 domain gates the barbed end and the efficiency with which its FH1 domain delivers profilin-actin to the barbed end [17, 25]. Formin isoforms have also been shown to possess specific nucleation activities, though the mechanism underlying these differences is not well understood [26, 27].

Actin polymerization proceeds at a rate that depends on the concentration of available actin monomers [2, 28]. This rate decreases as monomers are consumed over the course of polymerization. Because filament nucleation and elongation draw upon a common pool of actin monomers, both reactions contribute to the depletion of the monomer concentration and the rate at which one reaction proceeds influences the rate of the other. Modulation of the efficiency of one process can therefore alter the number of filaments assembled over the course of the reaction, as well as the lengths the filaments ultimately attain. Because formins mediate both nucleation and elongation, this interdependent mechanism might enable formin isoforms with differing polymerization activities to assemble filament networks with specific physical properties.

In this study, we dissected the mechanism of formin-mediated actin polymerization to determine the contributions of filament nucleation and elongation to the process of actin assembly. Using kinetic modeling, we found that the rates of both nucleation and elongation evolve over the course of a polymerization reaction. The period over which each process occurs is a key determinant of the number of filaments that are ultimately assembled, as well as their average equilibrium lengths. Inclusion of formin in polymerization reactions speeds filament nucleation, thus increasing the number and shortening the lengths of the filaments that are assembled. Analysis of *in vitro* polymerization assays confirmed the effects of varying the reactant concentrations and the filament elongation rate on polymerization. Although modulation of the elongation rate produces modest changes in the equilibrium lengths of formin-bound filaments, nucleation constitutes the primary mode of monomer consumption over the course of assembly. Sustained elongation of small numbers of formin-bound filaments therefore requires inhibition of nucleation via monomer sequestration and a low concentration of activated formin. Our results underscore the mechanistic advantage for keeping formin’s nucleation efficiency relatively low in physiological conditions [29]. This strategy also maximizes the impact of differences in the elongation properties of formin isoforms on the kinetics of actin network assembly.

## Materials and Methods

### Kinetic modeling

Kinetic schemes described by Sept and coworkers [2], Vavylonis and coworkers [18], and Paul and Pollard [10] were combined to generate a single model of actin polymerization that integrates the nucleation and elongation activities of formin in the absence and presence of profilin (Figure 1). Mathematical modeling of actin polymerization time courses was performed using COPASI [30] using previously published rate constants (Supplemental Table S1) [10, 18]. Time courses were calculated in deterministic mode using the LSODA algorithm [31, 32] at fixed initial bulk actin, profilin and/or formin concentrations.

**Figure 1.**
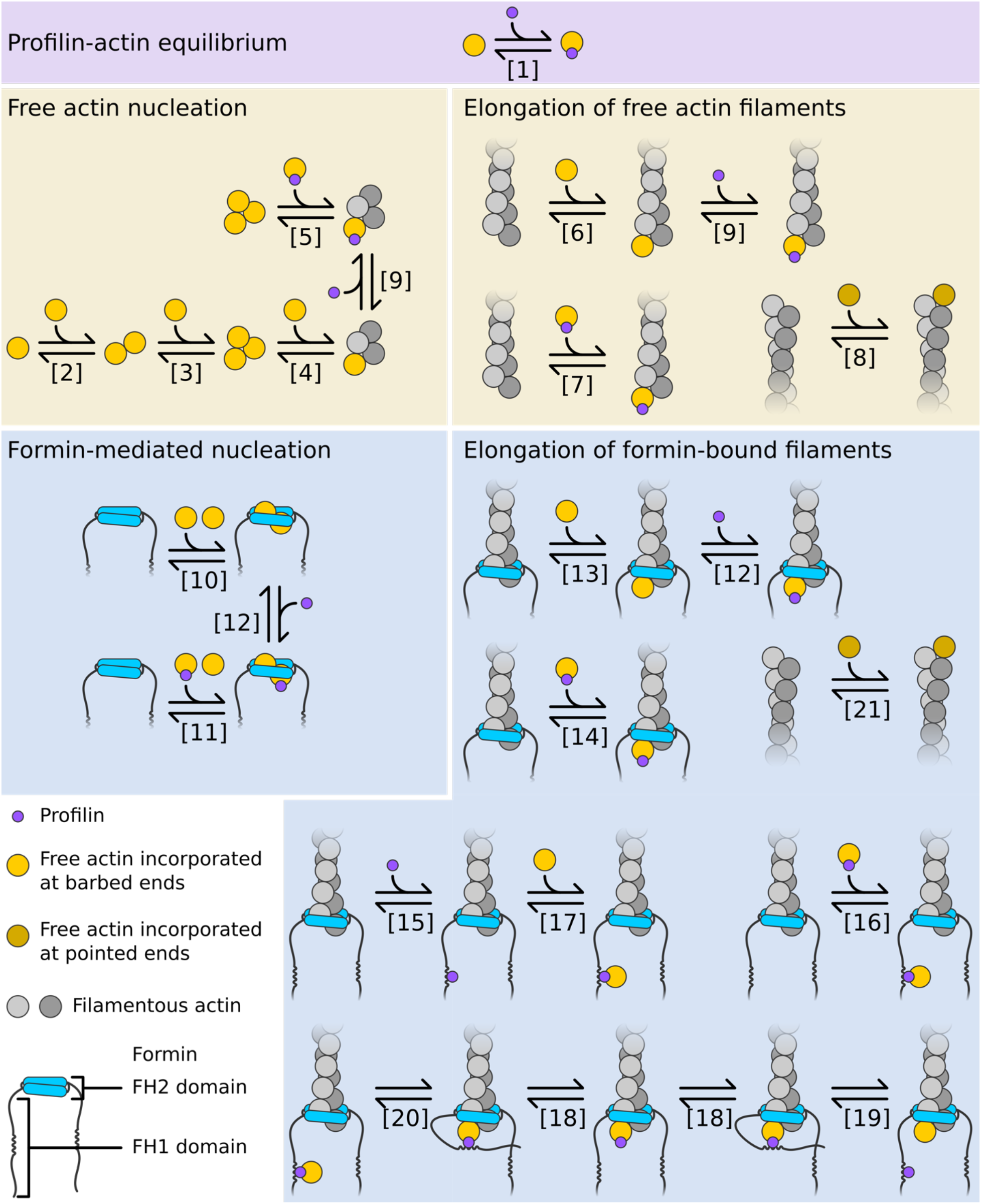
Kinetic model of spontaneous and formin-mediated actin filament nucleation and elongation. Profilin binds actin monomers (purple panel). Assembly of filaments with free barbed ends occurs via a nucleation phase and elongation at barbed and pointed ends (yellow panels). Assembly of filaments with formin-bound ends occurs via a nucleation phase, direct binding of actin and profilin-actin to barbed ends, FH1-mediated delivery of profilin-actin to the barbed end, and binding of actin monomers to pointed ends (blue panels). Rate constants corresponding to each reaction are described in Supplemental Table S1.

### Protein purification

Actin was purified from a chicken skeletal muscle acetone powder by one cycle of polymerization and depolymerization [33]. Monomeric actin was isolated by gel filtration on Sephacryl S-300 resin (GE Healthcare) in G-Buffer (2 mM Tris (pH 8.0), 0.2 mM ATP, 0.5 mM DTT, 0.1 mM CaCl_2_) and stored at 4°C. The concentration of actin was calculated using an extinction coefficient of 26,600 M^−1^cm^−1^ at 290 nm.

FH1FH2 constructs of Cdc12p (residues 882-1375) and Bni1p (residues 1227-1776) were expressed from pGEX-4T-3 plasmids (GE Healthcare) in BL21(DE3) RP Codon Plus cells and purified as previously described [34]. We used ProtParam (www.web.expasy.org/protparam [35]) to calculate extinction coefficients. *S. cerevisiae* profilin was expressed from a pMW172 vector in BL21 DE3 pLysS cells and purified as described [10, 34]. We used an extinction coefficient of 19,060 M^−1^cm^−1^ at λ = 280 nm to calculate the concentration of purified profilin.

### Microscopy and data analysis

Glass coverslips (22 mm x 50 mm; Fisher Scientific) were sonicated in 2% Hellmanex III (Millipore Sigma), rinsed extensively and sonicated in ddH_2_0. The imaging surface was constructed by placing Scotch Tape (3M) around the perimeter of a 4.5 mm x 4.5 mm region of the coverslip. Coverslips were flamed before use. The imaging surface was incubated with 0.5% Tween 20 in HS-TBS (600 mM NaCl, 50 mM Tris, pH 7.5) and 100 mg/mL bovine serum albumin (BSA) in HS-TBS. The imaging surface was washed in between each component and equilibrated with KMEI buffer (50 mM KCl, 1 mM MgCl_2_, 1 mM EGTA, 10 mM Imidazole (pH 7.0)) prior to introduction of the sample.

Ca^2+^-ATP-actin monomers were incubated with 0.05 mM MgCl_2_ and 0.2 mM EGTA for 3 minutes to generate Mg^2+^-ATP-actin. Polymerization of 2 µM actin monomers was initiated in the absence or presence of formin and/or profilin via incubation in KMEI buffer for 1-2 hours. Samples were taken every 30 minutes and analyzed via TIRF microscopy to determine when polymerization had reached equilibrium.

Assembled actin filaments were stabilized and fluorescently labeled via the addition of 4 µM fluorescein-isothiocyanate phalloidin (Sigma Aldrich). Following a 10-minute incubation, samples were diluted to a final concentration of 2-10 nM actin in 2x microscopy buffer (1x microscopy buffer: 10 mM Imidazole (pH 7.0), 50 mM KCl, 1 mM MgCl_2_, 1 mM EGTA, 50 mM DTT, 0.2 mM ATP, 15 mM glucose, 20 µg/mL catalase, 100 µg/mL glucose oxidase, 0.5 % (w/v) methylcellulose (4,000 cP at 2%)). Pipette tips were cut to reduce shearing of filaments during transfer, and 10-15 µL of the sample was loaded onto the imaging surface. The filaments were visualized by through-objective total internal reflection flurescence (TIRF) microscopy on an Olympus Ti83 motorized microscope equipped with a CellTIRF system using a 60x, 1.49 N.A. objective and a 488-nm laser. Images were acquired using a Hamamatsu C9100-23B ImagEM X2 EMCCD camera and CellSens Dimension software (Olympus).

Filament numbers and lengths were quantified from TIRF micrographs using a MATLAB program developed in-house. Filaments were detected from noise-filtered and background-subtracted images using MATLAB’s image thresholding algorithm. Detected filaments were skeletonized to facilitate length measurements. The number of filaments and their corresponding lengths were quantified for 3 replicates at each formin concentration using at least 5 fields of view per replicate. Single exponential fits were applied to filament length distributions [2]. For an exponential distribution, the fraction of filaments (*f_i_*) with length *l* was determined by the relation *f_i_* = λexp(-λ*l_i_*), where the mean length is 1/λ and the variance (*l_i_*) = (1/λ)^2^.

## Results

To dissect the contributions of filament nucleation and elongation to the dynamics of formin-mediated actin polymerization, we constructed a kinetic model composed of reaction schemes for both formin activities (Figure 1). Our model accounts for interactions among actin monomers, actin nuclei, filament barbed ends, filament pointed ends, formin and profilin. Consistent with published studies, we consider spontaneous filament assembly to occur through self-association of actin monomers into trimers [1, 2, 10]. Binding of a fourth monomer establishes a stable filament that can elongate via the association of additional, free actin monomers at both ends and profilin-bound monomers only at the barbed end [1, 28, 36–38]. Formin-mediated nucleation occurs via the association of a formin FH2 dimer with two actin monomers [9, 10]. Following this step, the formin remains bound to the barbed end of the filament [17, 39]. FH2-mediated gating slows barbed end elongation by decreasing the frequency of actin monomer binding [18, 19]. Profilin-actin complexes bind formin FH1 domains and are delivered to the barbed end in a diffusion-limited reaction [21, 22]. This process speeds elongation in a profilin-dependent manner [17, 18]. Formins do not influence pointed end elongation [17].

We used published values for the rate constants that govern each set of intermolecular associations (Supplemental Table S1) [10, 18]. Our model quantifies the number of filaments that are assembled over time and tracks subunit addition at filament ends. These measurements enable quantification of the nucleation and elongation rates as they evolve over the course of each simulated polymerization reaction.

### Actin polymerization in the absence of formin

To establish kinetic baselines for nucleation and elongation, we first simulated time courses of polymerization for reactions containing only actin monomers. Examination of 100,000 s trajectories enabled us to determine appropriate limits for making observations within the simulated framework. Consistent with published experimental observations [1], our simulations generated time courses of spontaneous actin polymerization that progress at rates that depend on the initial monomer concentration (Figure 2A). The total number of filaments assembled in each reaction increases over time and is largest at the highest sampled actin concentration at all time points throughout the course of the trajectory (Figure 2B). At actin concentrations exceeding 1 µM, polymerization trajectories reach stable values within approximately 15,000 s. Reactions containing less than 1 µM actin require more time to arrive at equilibrium.

**Figure 2.**
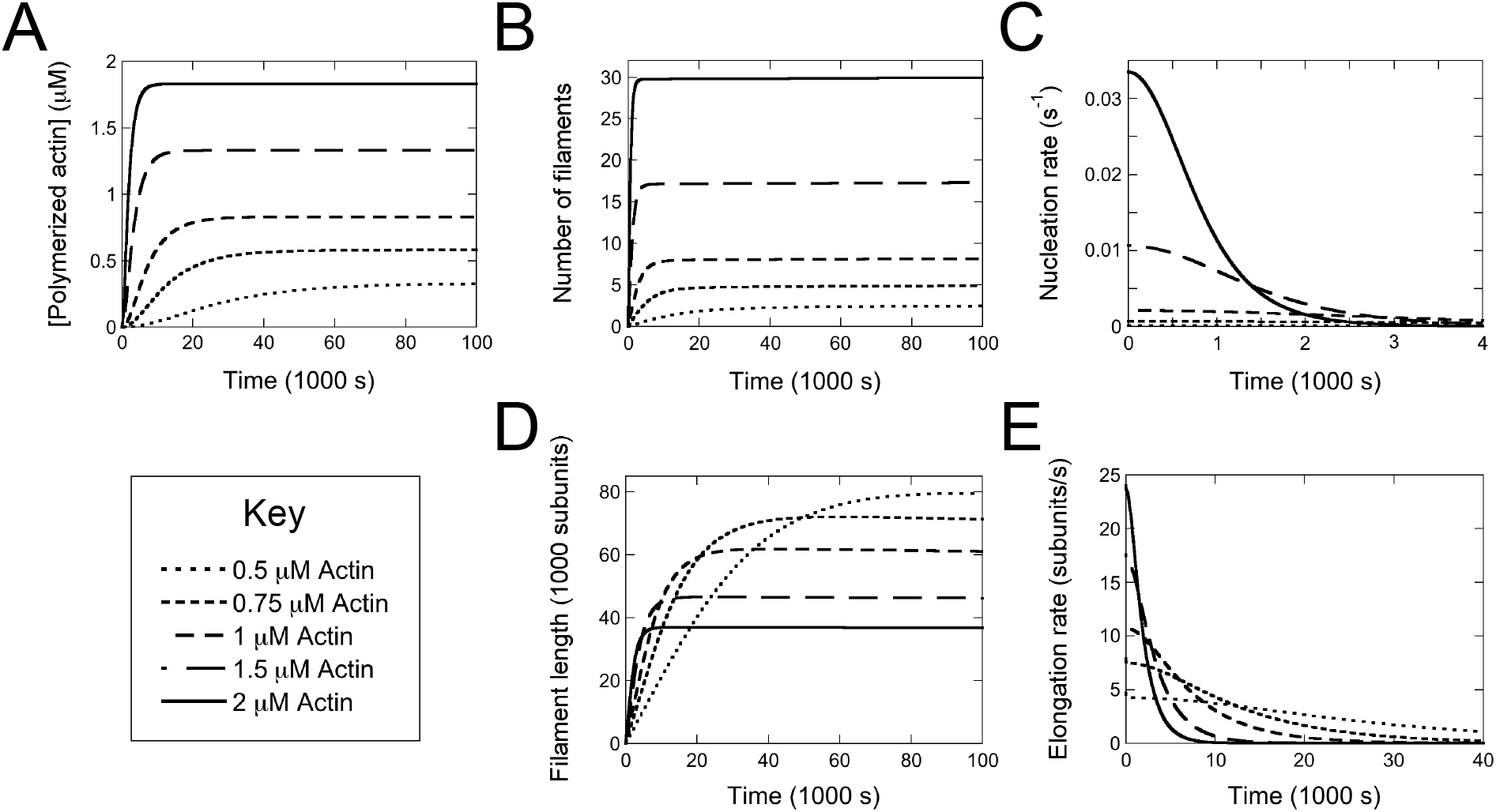
Actin polymerization in the absence of formin. Quantification of simulated polymerization reactions containing 0.5 µM (dotted line), 0.75 µM (short dashes), 1 µM (medium dashes), 1.5 µM (long dashes) and 2 µM (solid line) actin monomers. Each reaction was simulated over 100,000 s. (A) Concentration of polymerized actin over time. (B) Number of filaments assembled over time. (C) Actin filament nucleation rate over time. For clarity, data are plotted over a range of 4,000 s. (D) Average length of polymerized filaments over time. Lengths were calculated by dividing the concentration of polymerized actin by the number of filaments at each time point. (E) Rate of filament elongation over time. For clarity, data are plotted over a range of 40,000 s.

As expected, filament nucleation rates are fastest at early time points and decrease over time as monomers are consumed by polymerization (Figure 2C). The initial nucleation rate is faster at high actin concentrations than at low concentrations. However, the shape of the nucleation trajectory broadens in reactions containing lower actin concentrations, leading to faster nucleation rates at later points in the trajectory. At the lowest actin concentrations we simulated, the nucleation rate trajectory is essentially flat, but its value overtakes the nucleation rates of the higher concentrations as the reaction progresses.

In each polymerization reaction, filament elongation takes place over a longer period than nucleation (Figure 2E). The initial filament elongation rate is fastest at the highest actin concentration. However, reactions containing lower actin concentrations exhibit broadened trajectories with faster elongation rates at later times. Thus, a decrease in the rate at which monomers are consumed prolongs the time over which individual filaments elongate through monomer binding. As a result, filament lengths measured at the ends of the simulated trajectories are inversely proportional to the initial concentration of monomeric actin (Figure 2D).

To assess the relevance of our kinetic modeling to experimental observations, we compared our simulated polymerization reactions to *in vitro* measurements of assembled filaments. We incubated purified actin monomers in conditions mimicking those in our simulated reactions. Once the reactions reached equilibrium, we added fluorescent phalloidin, imaged the filaments using total internal reflection fluorescence (TIRF) microscopy and quantified filament numbers and lengths.

In addition to nucleation and elongation, actin filaments assembled *in vitro* undergo length-dependent fragmentation and annealing [2, 40]. These reactions can alter both the number and lengths of the actin filaments. The probability of severing and annealing both increase with the concentration of filaments [2]. To minimize the likelihood of these events occurring, we used actin monomer concentrations in the low micromolar range.

Each *in vitro* reaction robustly assembled into filaments of varying lengths (Figure 3A). As previously reported [2], the distributions of filament lengths are well characterized by single exponential fits (Figure 3B), which yield both an average filament length and a variance (see Methods). The average lengths of filaments assembled in our reactions containing 2 µM actin agree with published measurements performed on similar reactions, confirming that our sample preparation and visualization methods minimize filament breakage [2]. As the concentration of actin monomers included in each reaction increases, the number of assembled filaments increases and the average filament length decreases (Figure 3C and D; data points). The magnitude of these concentration-dependent changes is similar to the trend predicted by our simulations (Figure 3C and D; lines), confirming that our model produces physiologically relevant insights at these actin concentrations.

**Figure 3.**
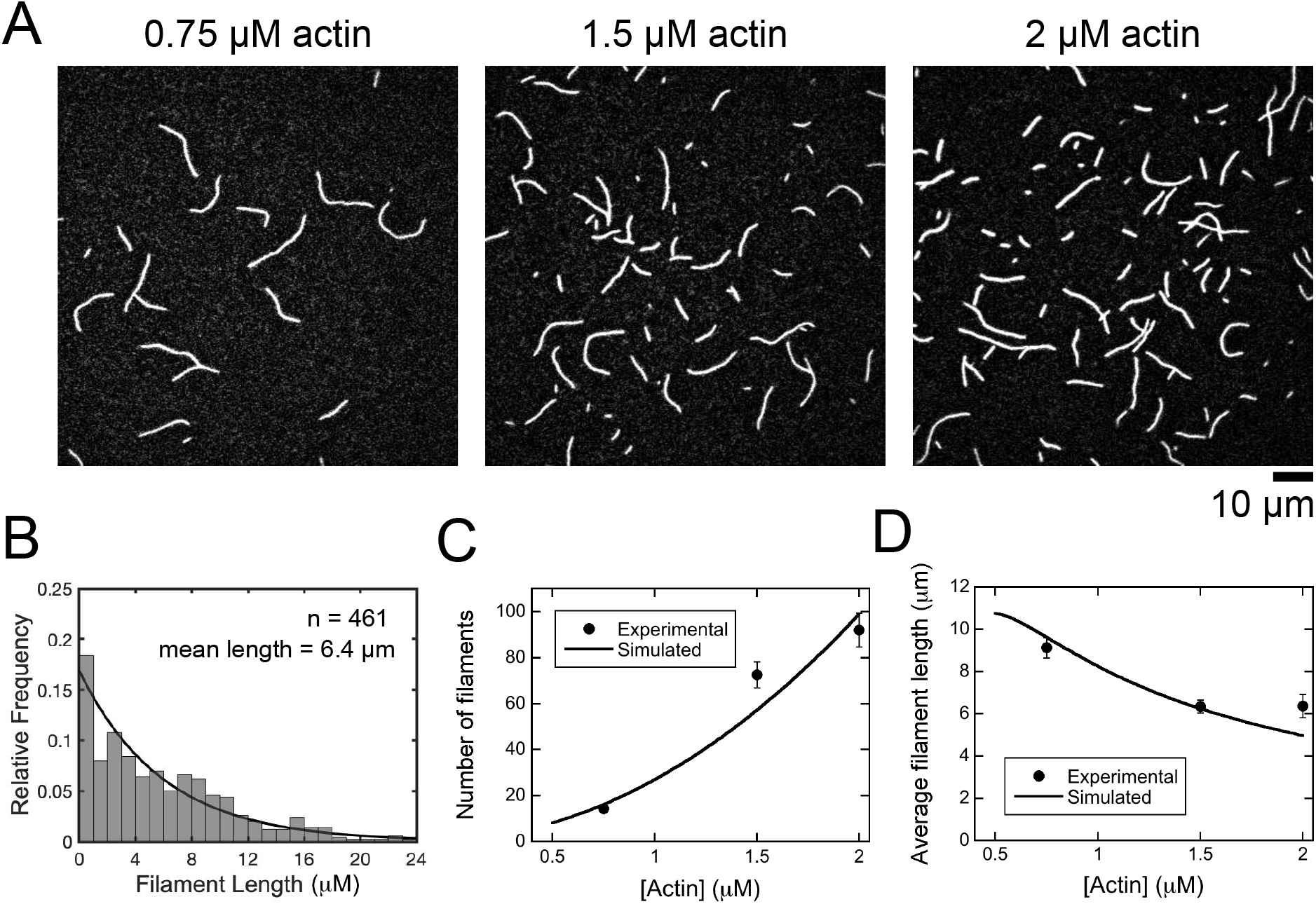
*In vitro* actin polymerization in the absence of formin. The experimental conditions were as follows: A range of concentrations of actin monomers in polymerization buffer. The filaments were labeled with Alexa 488-phalloidin and visualized by TIRF microscopy. (A) Representative TIRF micrographs of filaments assembled in reactions containing a range of actin concentrations. (B) Histogram of filament lengths measured at equilibrium for a representative polymerization reaction containing 2 µM actin. The line is an exponential fit to the data. The fitted value for λ is 0.156, and the mean filament length is 1/λ, or 6.4 µm. (C) Dependence of the number of actin filaments visualized per 10,000 µm^2^ on the actin concentration. Error bars are standard errors of the mean of at least 5 micrographs. Simulated data (solid line) were normalized and plotted on the same y-axis scale as the experimental data. (D) Dependence of the average filament length on the actin concentration. Error bars are standard errors of the mean of at least 5 micrographs. Simulated data (solid line) were normalized and plotted on the same y-axis scale as the experimental data.

### Formins shorten the time frame for nucleation

Our simulated trajectories indicate that varying the concentration of the components in a polymerization reaction can alter the number of filaments that are assembled, as well as their lengths at equilibrium. To investigate how the nucleation and elongation activities of formins influence the distribution of actin monomers into populations of filaments, we simulated actin polymerization in the presence of formin. We initially sampled a range of formin concentrations spanning 6 orders of magnitude. In the absence of profilin, FH2 domain gating limits the rate at which actin monomers bind filament barbed ends, thus slowing the elongation of formin-bound filaments relative to filaments with free barbed ends [17]. To create distinct rates of elongation for our simulated formin-bound and free filaments, we used an FH2 gating factor of 0.5. To ensure that our reaction time courses arrive at equilibrium within a reasonable time frame, we performed our simulations with 2 µM actin monomers. Under these conditions, most of our simulated trajectories reach equilibrium within 10,000 s (∼2.8 hours).

In each simulated reaction, the number of assembled filaments increases over time, and reactions that include larger concentrations of formin contain more filaments at all time points than reactions that include lower concentrations of formin (Figure 4A). The initial formin-mediated nucleation rate is linearly proportional to the concentration of formin, consistent with a nucleation reaction that involves the association of two actin monomers with one pre-assembled, stable formin dimer (Figure 4B, y-intercept values). In contrast, nucleation of filaments with free barbed ends (i.e., not formin-bound) occurs at the same initial rate independent of the formin concentration (Supplemental Figure S1A). The formin-mediated and spontaneous nucleation rates both decrease over time, and the time period over which both types of nucleation events occur depends on the formin concentration (Figure 4B and Supplemental Figure S1A). At each formin concentration we sampled, formin-mediated nucleation occurs over a period that is approximately twice as long as the time period for spontaneous nucleation.

**Figure 4.**
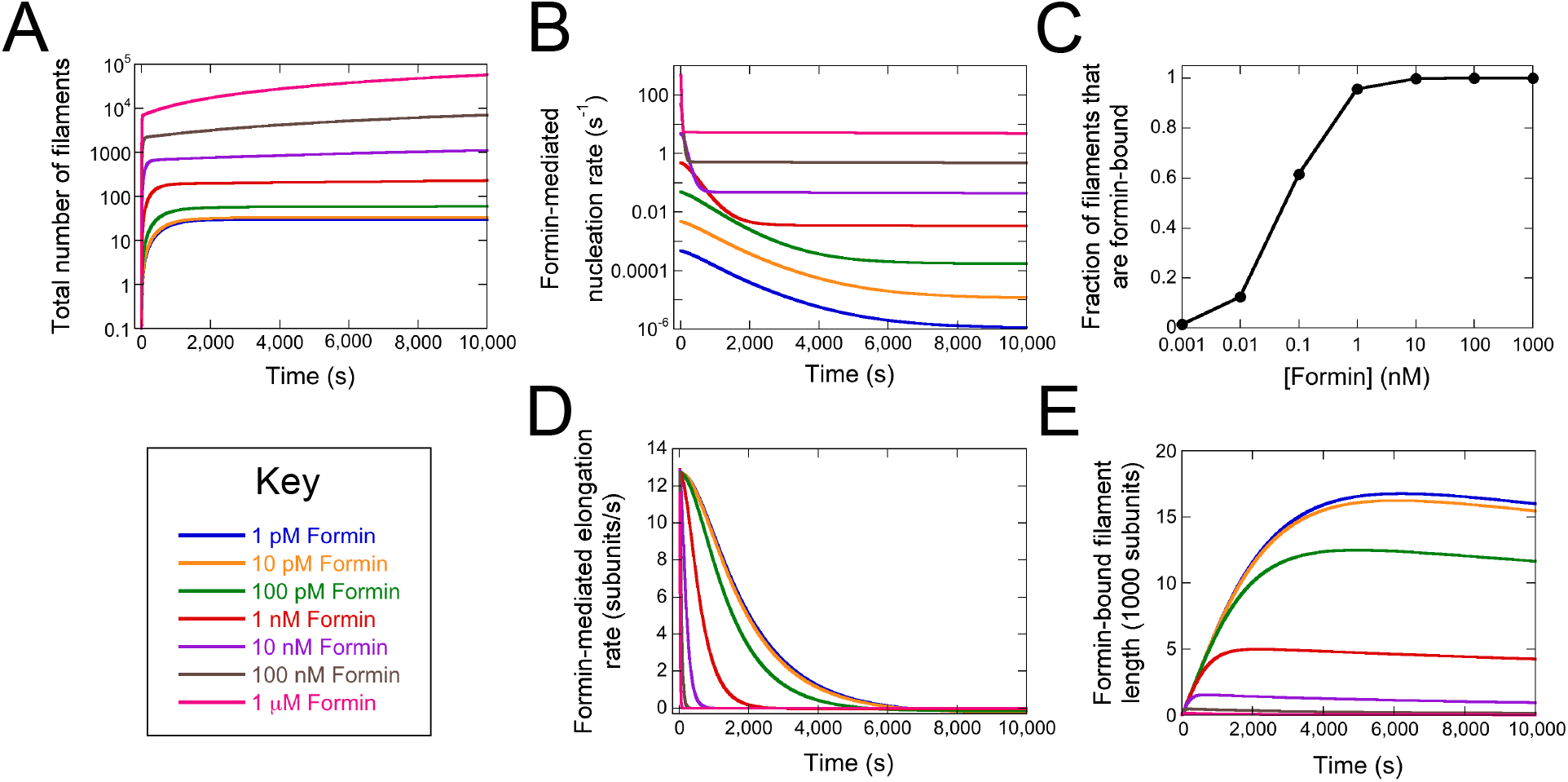
Formins nucleate short filaments. Quantification of simulated polymerization reactions containing 2 µM actin and 1 pM (blue lines), 10 pM (orange lines), 100 pM (green lines), 1 nM (red lines), 10 nM (purple lines), 100 nM (brown lines) or 1 µM (pink lines) formin. The formin gating factor was set at 0.5. Each simulation was carried out over 10,000 s. (A) Total number of filaments assembled over the course of each trajectory. The total number of filaments was determined by summing the numbers of formin-bound and free filaments at each time point. For clarity, the y-axis is represented on a log scale. (B) Formin-mediated nucleation rates over time. For clarity, the y-axis is represented on a log scale. (C) The dependence of the fraction of filaments that are formin-bound on the concentration of formin. For clarity, the x-axis is represented on a log scale. (D) Formin-mediated elongation rate over time. (E) Average lengths of formin-bound filaments over time.

The equilibrium composition of each polymerized reaction depends on the formin concentration. At 1 µM formin, the rate of formin-mediated nucleation is ∼10^5^ times faster than nucleation of filaments with free barbed ends (Figure 4B and Supplemental Figure S1A). Under these conditions, the polymerized reaction contains mostly formin-bound filaments (Figure 4C). In contrast, spontaneous actin nucleation is faster than formin-mediated nucleation at 1 pM formin. Thus, the majority of filaments assembled in this reaction have free barbed ends. At intermediate concentrations of formin, the rate of formin-mediated filament nucleation approaches the spontaneous nucleation rate. As a result, polymerized reactions contain mixtures of both formin-bound and free filaments.

### Formins assemble short filaments

Nucleated filaments elongate at rates that depend on whether their barbed ends are formin-bound or free [17]. In our simulated reactions, elongation of both formin-bound and free filaments slows over time as monomers are consumed (Figure 4D and Supplemental Figure S1B). The elongation rate trajectory narrows as the formin concentration increases and reactions containing lower concentrations of formin exhibit faster elongation rates at later times. As a result, the average filament length quantified once each reaction has attained equilibrium is inversely dependent on the formin concentration (Figure 4E and Supplemental Figure S1C).

Consistent with our kinetic modeling, inclusion of an FH1FH2 construct of the *S. cerevisiae* formin Bni1p (which has a gating factor of 0.5 [10, 17]) increases the number of filaments assembled in polymerization reactions performed *in vitro* (Figure 5A, top row; and 5B, filled circles). The observed increase in filament nucleation is matched by a decrease in the average filament length (Figure 5C, filled circles). Varying the concentration of Bni1p produces hyperbolic effects on both filament number and length that reach a plateau at concentrations above 500 nM Bni1p. Although our model is not specifically tailored to Bni1p, these *in vitro* measurements phenomenologically reproduce the trends predicted by our simulations (Figure 5D and E; gating = 0.5).

**Figure 5.**
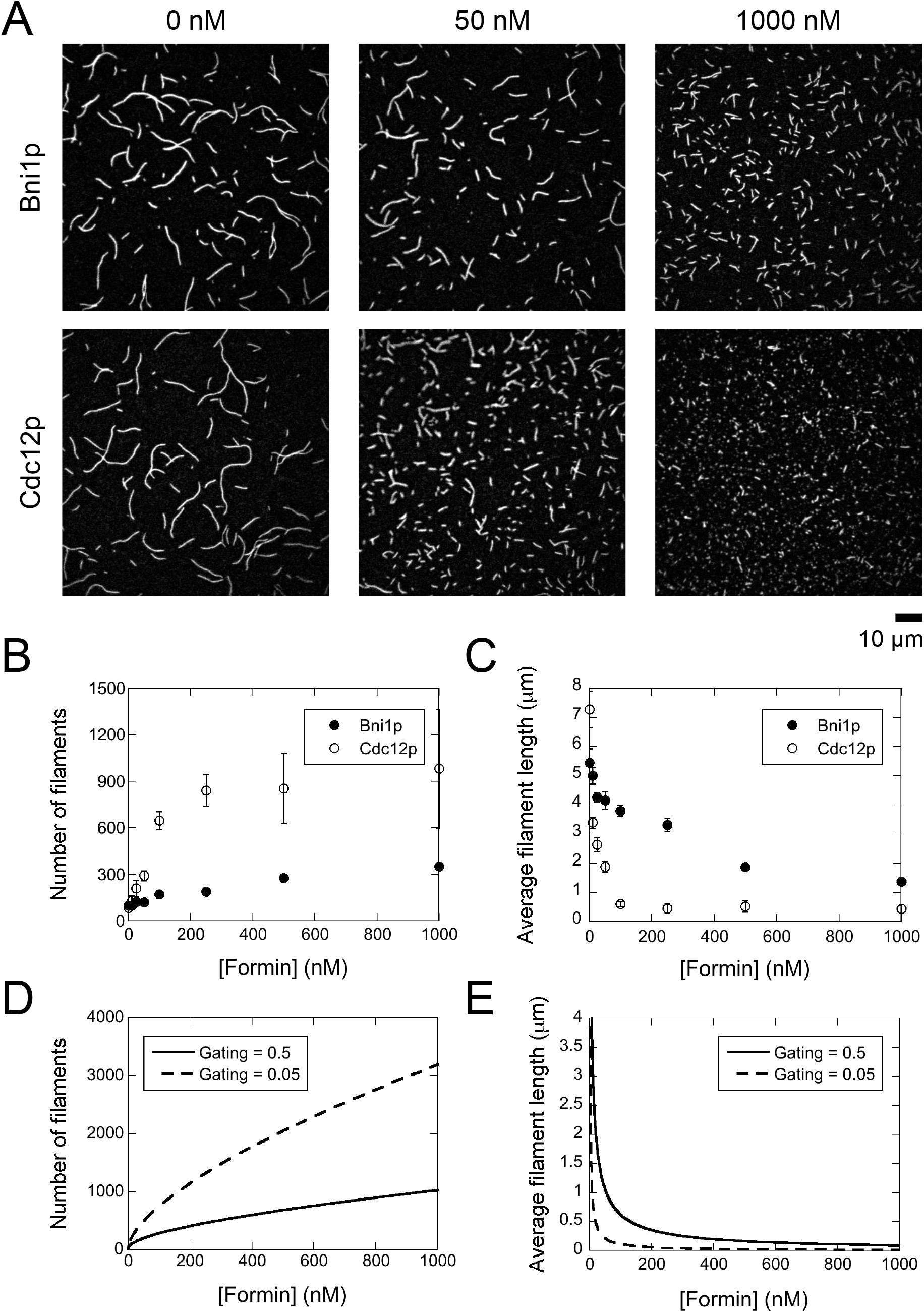
*In vitro* formin-mediated actin polymerization. The experimental conditions were as follows: 2 µM actin monomers and a range of concentrations of FH1FH2 constructs of Bni1p or Cdc12p in polymerization buffer. The filaments were labeled with Alexa 488-phalloidin and visualized by TIRF microscopy. (A) Representative TIRF micrographs of filaments assembled in the absence and presence of Bni1p (top row) or Cdc12p (bottom row). (B) Dependence of the number of actin filaments visualized per 10,000 µm^2^ on the concentration of Bni1p (filled circles) or Cdc12p (open circles). Error bars are standard errors of the mean of at least 5 micrographs. (C) Dependence of the average filament length on the concentration of Bni1p (filled circles) or Cdc12p (open circles). Error bars are standard errors of the mean of at least 5 micrographs. (D) Simulated dependence of the number of filaments on the concentration of formin with a gating factor of 0.5 (solid line) or 0.05 (dashed line). (E) Simulated dependence of the average filament length on the concentration of formin with a gating factor of 0.5 (solid line) or 0.05 (dashed line).

### Filament length distributions depend on FH2 domain gating

Our simulations demonstrate that inclusion of a formin that both stimulates nucleation and slows elongation increases the number and shortens the lengths of actin filaments assembled over the course of polymerization. To assay the dependence of filament lengths on the rate of elongation, we varied the gating factor of our simulated formin. The gating factor “p” is a measure of the probability that a formin FH2 dimer adopts a polymerization-competent conformation [18]. Formins with gating factors near 0 inhibit subunit addition at barbed ends nearly completely. In contrast, barbed ends bound by formins with a gating factor of 1 elongate at the same rate as do filaments with free barbed ends. To simplify our analysis, we performed our simulations at a formin concentration of 1 nM, which ensures that over 95% of filaments assembled over the course of each trajectory are formin-bound.

At each gating factor we sampled, formin-mediated actin assembly reaches equilibrium within 5,000 s. Filaments nucleated by formin elongate over time and attain lengths that depend non-linearly on the gating factor (Figure 6A). Filament length is most sensitive to changes in gating when the gating factor is small. For example, decreasing the gating factor from 0.5 to 0.1 decreases the average filament length by ∼55%. In contrast, increasing the gating factor from 0.5 to 0.9 increases the average length by only ∼33%.

**Figure 6.**
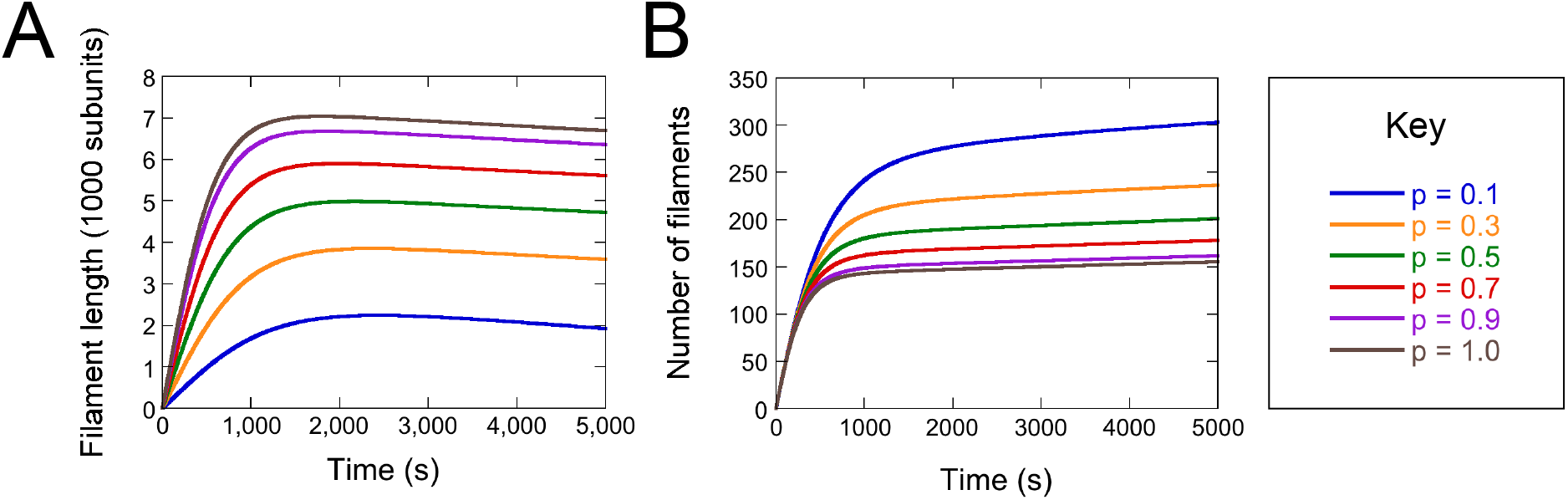
Filament length and number depend nonlinearly on FH2 domain gating. Quantification of simulated polymerization reactions containing 2 µM actin and 1 nM formin at a range of gating factors. The gating factor “p” values were 0.1 (blue lines), 0.3 (orange lines), 0.5 (green lines), 0.7 (red lines), 0.9 (purple lines) and 1.0 (brown lines). Each simulation was carried out over 10,000 s. (A) Average filament length over time. (B) Number of filaments assembled in each reaction over time.

The number of filaments assembled in each polymerization reaction also depends non-linearly on the gating factor (Figure 6B). A formin with a gating factor of 0.1 nucleates 50% more filaments than a formin with a gating factor of 0.5, which in turn nucleates ∼33% more filaments than a formin with a gating factor of 0.9.

To assess the consequences of varying the gating factor, we repeated our *in vitro* actin polymerization reactions in the presence of a range of concentrations of the *S. pombe* formin Cdc12p (Figure 5A, bottom row). This formin has a gating factor of ∼0.05 and therefore filaments nucleated by this formin elongate mainly through pointed end elongation [17]. Inclusion of Cdc12p stimulates the assembly of shorter and more numerous filaments than observed in reactions containing identical concentrations of Bni1p (Figure 5B and C). Although we observe fewer and longer filaments *in vitro* than in our simulations, the magnitude of the differences in both the number and the lengths of the filaments assembled by each formin closely resembles the differences predicted by our simulations (see Discussion) (Figure 5D and E).

### Inhibition of nucleation by profilin promotes the assembly of long filaments

Formins overcome the limitations on elongation imposed by FH2 gating by binding and delivering profilin-actin complexes to filament barbed ends via their FH1 domains [18]. In addition to speeding formin-mediated elongation, profilin also sequesters monomers and inhibits both nucleation and pointed end elongation [38, 41, 42]. It is therefore likely that profilin influences the distribution of actin monomers into polymerized filaments in both the absence and presence of formin.

To obtain baseline measurements for the effects of profilin on polymerization, we simulated the effects of a range of profilin concentrations on actin assembly in the absence of formin. We set the affinity at which profilin binds to actin monomers to 3 µM, which matches the measured binding constant for *S. cerevisiae* profilin [43]. Consistent with its role as a monomer sequestration protein, we found that profilin decreases the initial nucleation rate relative to the rate observed for actin alone (Figure 7A). The magnitude of this effect is proportional to the fraction of actin monomers that are profilin-bound in each reaction. Reactions containing at least 10 µM profilin undergo minimal nucleation owing to the near-complete binding of actin monomers to profilin. Concomitantly, the time period over which nucleation occurs increases as the concentration of profilin increases.

**Figure 7.**
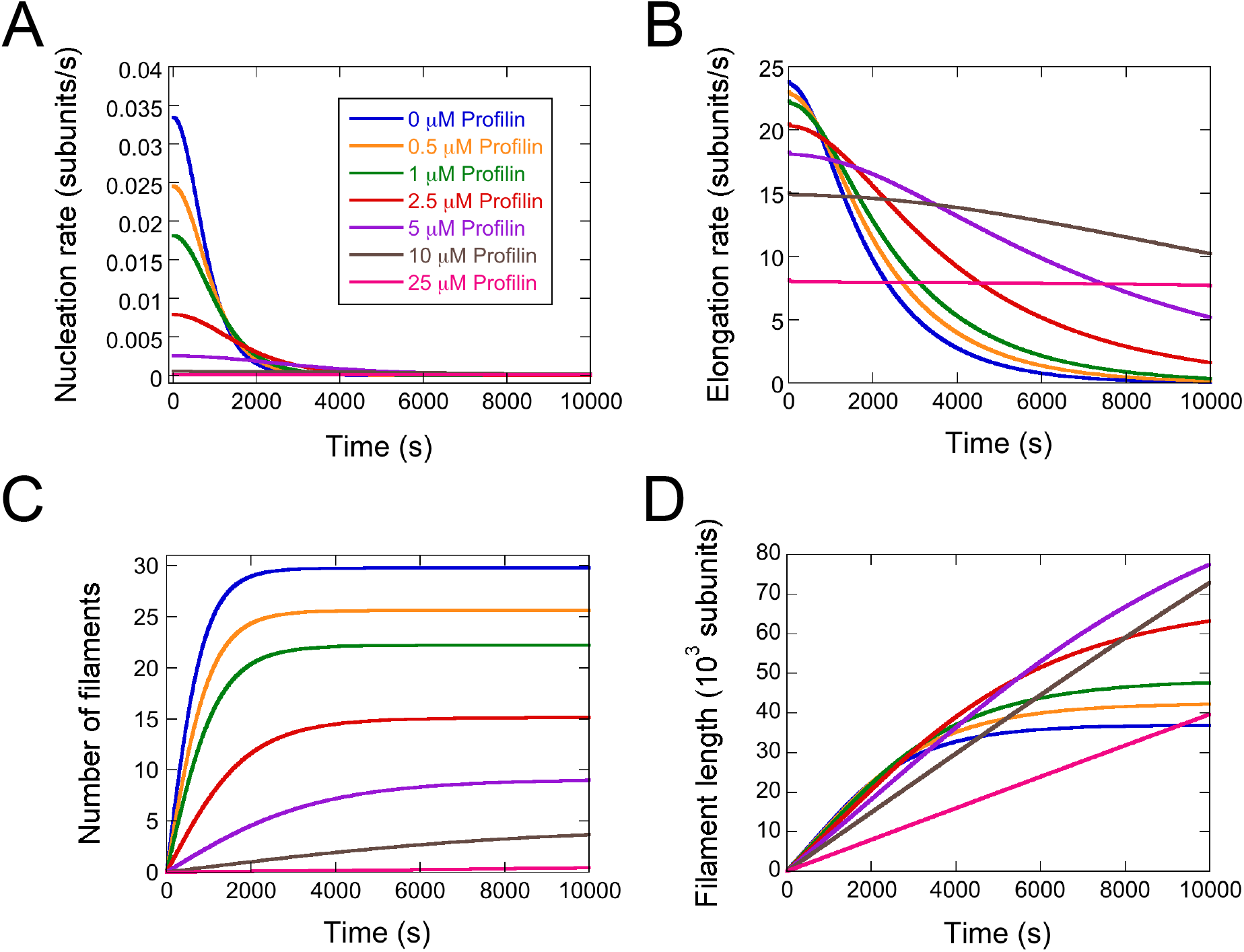
Inhibition of nucleation by profilin promotes the assembly of long filaments. Quantification of simulated polymerization reactions containing 2 µM actin and 0 µM (blue lines), 0.5 µM (orange lines), 1 µM (green lines), 2.5 µM (red lines), 5 µM (purple lines), 10 µM (brown lines) or 25 µM (pink lines) profilin. Profilin’s affinity for actin monomers was set at 3 µM. Each simulation was carried out over 10,000 s. (A) Filament nucleation rate over time. (B) Filament elongation rates over time. (C) Number of filaments assembled in each reaction over time. (D) Average filament length over time.

The initial rate of filament elongation also slows in the presence of profilin, but this effect is less sensitive to the profilin concentration compared to the nucleation rate (Figure 7B). Unlike nucleation, which depends on the concentration of free (i.e., not profilin-bound) monomers, filament elongation is limited by the dissociation of profilin from the barbed end [44, 45]. Profilin’s affinity for barbed ends is 100-fold weaker than its affinity for monomers [37]. Thus, the weaker dependence of the initial elongation rate on the profilin concentration is consistent with this difference in affinity. In contrast to the modest effect on the initial rate of filament elongation, profilin significantly increases the time over which filament elongation proceeds. As a result, filaments elongate at near-constant rates for at least 5,000 s in reactions containing at least 10 µM profilin. These reactions also require longer than 100,000 s simulations to reach equilibrium. In combination, profilin’s effects on nucleation and elongation result in a concentration-dependent decrease in the number of assembled actin filaments, as well as an increase in filament length (Figure 7C and D).

### Profilin modulates the lengths of filaments assembled by formin

Inclusion of 1 nM formin in polymerization reactions containing profilin significantly increases the initial rate of filament nucleation compared to reactions conducted in the absence of formin (Figure 8A). As the profilin concentration increases, the magnitude of formin’s stimulatory effect on the initial nucleation rate increases (Figure 8B). Formin also shortens the period over which nucleation occurs at all concentrations of profilin, and all reactions come to completion within 1500 s (Figure 8A).

**Figure 8.**
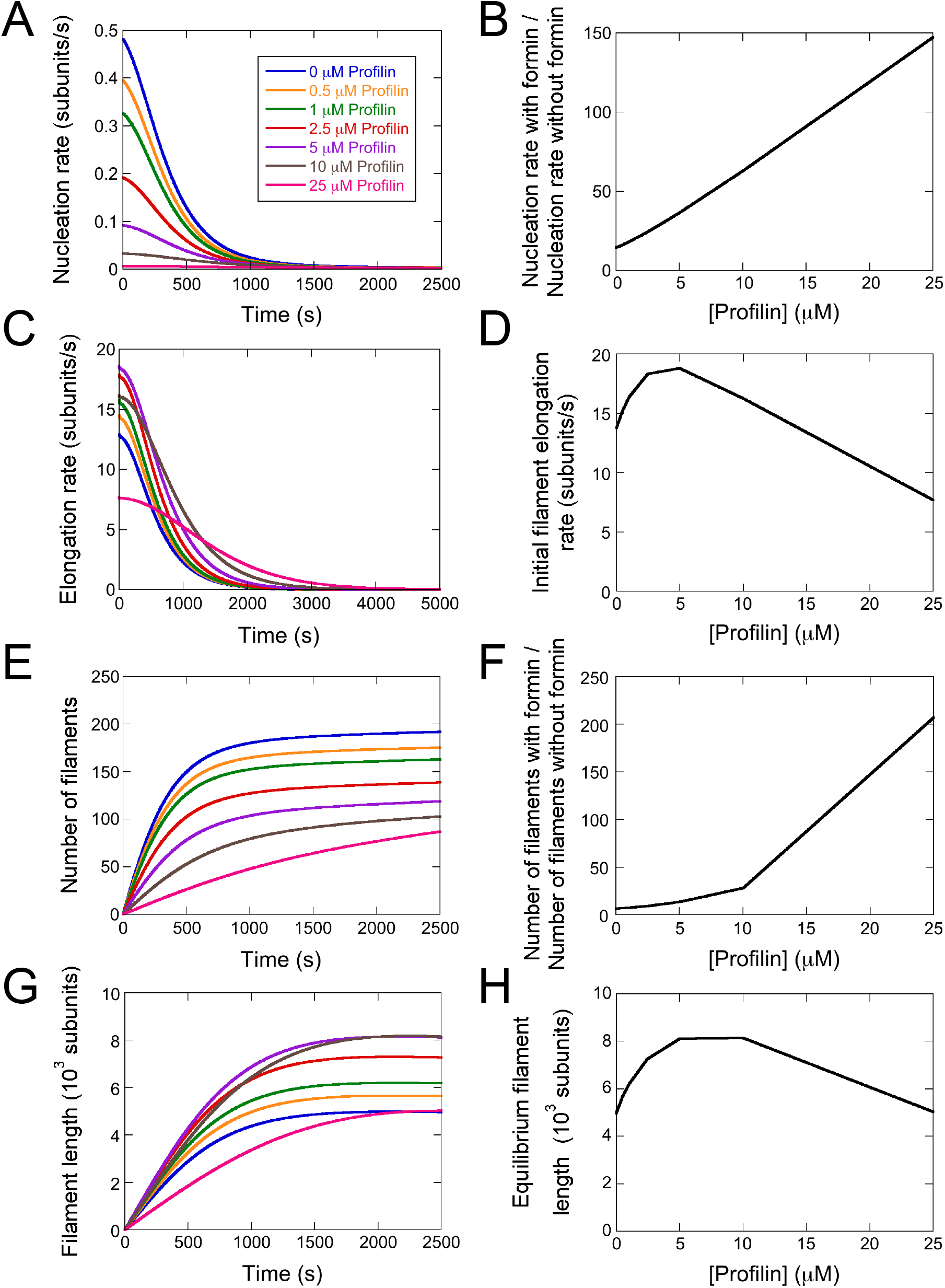
Profilin modulates the lengths of filaments assembled by formins. Quantification of simulated polymerization reactions containing 2 µM actin, 1 nM formin and 0 µM (blue lines), 0.5 µM (orange lines), 1 µM (green lines), 2.5 µM (red lines), 5 µM (purple lines), 10 µM (brown lines) or 25 µM (pink lines) profilin. Profilin’s affinity for actin monomers was set at 3 µM. Each simulation was carried out over 10,000 s. (A) Filament nucleation rate over time. (B) Dependence of the ratio of the initial nucleation rate measured in the presence of formin to the rate measured in the absence of formin on the concentration of profilin. (C) Filament elongation rates over time. (D) Dependence of the initial filament elongation rate (corresponding to the y-intercept in panel (C)) on the concentration of profilin. (E) Number of filaments assembled in each reaction over time. (F) Dependence of the ratio of the number of filaments assembled in the presence of formin to the number measured in the absence of formin on the concentration of profilin. Filament numbers were quantified once each reaction reached equilibrium. (G) Average filament length over time. (H) Dependence of the average equilibrium filament lengths on the concentration of profilin.

Profilin produces a well-characterized biphasic change in the rate of formin-mediated elongation [10, 17, 34]. Sub-saturating concentrations of profilin speed elongation by producing profilin-actin complexes that bind formin FH1 domains, enabling their delivery to the barbed end. At concentration of profilin that exceed the concentration of actin monomers, competition among profilin-actin complexes and free profilin for binding to FH1 domains slows filament elongation [18]. Consistent with these established effects on elongation, the initial elongation rate in simulations that include formin depends non-linearly on the profilin concentration and is fastest at 5 µM profilin (Figure 8C and D). Elongation slows over time, and the period over which elongation proceeds increases as a function of profilin (Figure 8C). At all concentrations of profilin, elongation takes place over a longer period of time than does nucleation.

Together, the effects of profilin on filament nucleation and elongation decrease the total number of filaments assembled over the course of a polymerization reaction (Figure 8E). Despite this effect, all reactions containing formin nucleate at least six times as many filaments as the same reaction conducted in the absence of formin (Figures 7C and 8F). Formin’s relative effect on the number of assembled filaments increases with increasing profilin concentration. Filaments assembled in reactions containing formin also attain their maximum lengths much faster than filaments assembled in the absence of formin (Figures 7D and 8G). These filament lengths are approximately 10 times shorter than filaments polymerized in the absence of formin, consistent with the significantly larger number of filament ends generated by nucleation across which to distribute actin monomers. Reactions containing 5 and 10 µM profilin produce the longest filaments, whereas filaments are shortest in reactions containing 0 and 25 µM profilin (Figure 8H). These equilibrium filament lengths generally mirror the effects of profilin on the rate of formin-mediated elongation, but the trend is shifted to slightly higher profilin concentrations. This shift likely results from profilin’s own effects on actin polymerization via its role as a monomer-sequestering protein.

### In vitro polymerization reactions capture the effects of formin and profilin on filament assembly

We compared our simulated results to measurements of actin assembly reactions performed *in vitro* in the presence of 5 µM profilin. We introduced a range of concentrations of our FH1FH2 construct of Bni1p and quantified the number of filaments and their average lengths once each reaction reached equilibrium (Figure 9A). Bni1p FH1FH2 produces a concentration-dependent increase in the number of filaments assembled in our reactions (Figure 8B; filled circles). This increase in filament number is matched by a shortening of the average filament length (Figure 9C). Both effects phenomenologically reproduce the trends predicted by our simulations (Figure 9D and E). In reactions containing at least 250 nM formin, assembled filaments are approximately twice as long as filaments polymerized in reactions containing the identical concentration of Bni1p but lacking profilin (Figures 5C and 9C).

**Figure 9.**
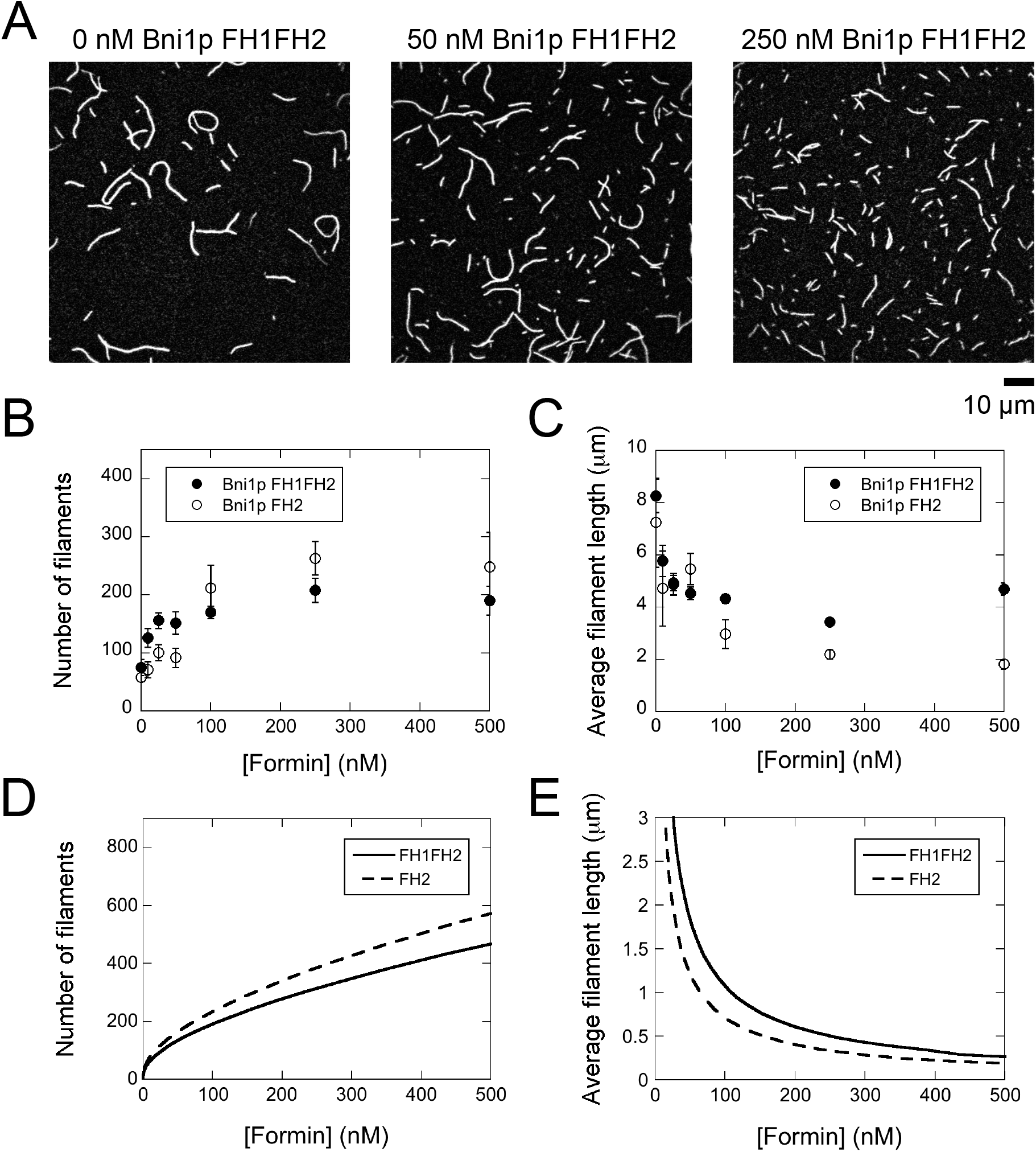
*In vitro* polymerization reactions capture the effects of formin and profilin on filament assembly. The experimental conditions were as follows: 2 µM actin monomers, 5 µM *S. cerevisiae* profilin and a range of concentrations of Bni1p FH1FH2 or FH2 in polymerization buffer. The filaments were labeled with Alexa 488-phalloidin and visualized by TIRF microscopy. (A) Representative TIRF micrographs of filaments assembled in the absence and presence of Bni1p FH1FH2. (B) Dependence of the number of actin filaments visualized per 100 µm^2^ on the concentration of Bni1p FH1FH2 (filled circles) or Bni1p FH2 (open circles). Error bars are standard errors of the mean of at least 5 micrographs. (C) Dependence of the average filament length on the concentration of Bni1p FH1FH2 (filled circles) or FH2 (open circles). Error bars are standard errors of the mean of at least 5 micrographs. (D) Dependence of the number of filaments on the concentration of FH1FH2 (solid line) or FH2 (dashed line) in simulated reactions. (E) Dependence of the average filament length on the concentration of FH1FH2 (solid line) or FH2 (dashed line) in simulated reactions.

To test the relationship between the FH1-mediated filament elongation rate and filament lengths measured at equilibrium, we compared the results of our experiments conducted with Bni1p FH1FH2 to measurements performed in the presence of Bni1p FH2. Because the FH2 construct lacks profilin-binding sites, it mediates slower filament elongation than does the FH1FH2 construct in the presence of profilin [34]. Use of this construct therefore enabled us to modulate the formin-mediated elongation rate while keeping the profilin concentration constant at 5 µM.

Consistent with its strong nucleation activity, Bni1p FH2 produces a concentration-dependent increase in the number of filaments assembled in each polymerization reaction, as well as a decrease in the average filament length (Figure 9B and 9C, open circles). At most formin concentrations, Bni1p FH2 assembles shorter filaments than does Bni1p FH1FH2. The difference in the average lengths of filaments polymerized by these two Bni1p constructs is smaller than the difference in their elongation rates but is consistent with the trend predicted by our simulations (Figure 9E) [34]. Bni1p FH2 also nucleates more filaments than does Bni1p FH1FH2 at most formin concentrations (Figure 9B and 9D). This suggests that a slower rate of filament elongation boosts the efficiency of filament nucleation even for formin constructs with identical FH2 domains. This increase in nucleation stimulates *de novo* filament assembly at the expense of filament length.

## Discussion

Formins regulate actin filament nucleation and elongation in a profilin-dependent manner. Both processes require and consume actin monomers, suggesting that the nucleation and elongation activities of formins might be interdependent. To dissect the contributions of the nucleation and elongation reactions to formin-mediated actin assembly, we constructed a kinetic model that enables independent examination of each process throughout polymerization. We found that the rates of nucleation and elongation decrease over the course of polymerization and that changes in these rates alter the number and lengths of the resulting actin filaments.

### Filament elongation occurs over a longer time period than nucleation

Formins mediate actin polymerization at rates that depend on the concentration of actin, formin and profilin. To assess the role of each component in actin assembly, we considered each protein individually and in combination. In all of our simulated reactions, the rates of nucleation and elongation are fastest at initial timepoints and decrease over the course of the reaction. In most reactions, elongation occurs over a longer period than does nucleation, indicating that monomer addition at pre-existing filament ends occurs at low actin concentrations that do not freely support the formation of nuclei.

In reactions containing actin alone, variation of the actin concentration produces changes in the equilibrium lengths of the assembled filaments. Filament lengths become shorter as the concentration of actin increases, consistent with a nonlinear increase in the number of filament ends. This nonlinear relationship arises from the dependence of the nucleation rate on the cube of the monomer concentration (Figure 1) [2]. An increase in the amount of available actin therefore produces a large increase in the number of assembled filaments. These new filaments in turn increase the number of binding sites across which the monomer pool is distributed via elongation. As a result, each filament incorporates fewer actin monomers and ultimately attains a shorter length at equilibrium.

Inhibition of nucleation via profilin-mediated monomer sequestration decreases the number of filaments that are produced over the course of each reaction. The magnitude of this decrease depends on the fraction of monomers that are profilin-bound, which is dictated by profilin’s affinity for monomers. Once nucleated, filaments can bind profilin-actin complexes at their barbed end. Dissociation of profilin following each profilin-actin binding event regenerates the barbed end binding site and enables sustained filament elongation. Although profilin’s affinity for barbed ends is ∼100-fold weaker than its affinity for monomers [37], this dissociation step becomes slower as the profilin concentration increases [44, 45]. In combination, these effects on nucleation and elongation dramatically slow polymerization and produce populations of filaments that are smaller in number and attain longer average lengths as the concentration of profilin increases.

In contrast to the effects of profilin, inclusion of formin promotes filament nucleation at the expense of elongation. However, variation of the elongation rate can also modulate the observed filament nucleation rate. Varying the formin gating factor alters the period of time over which both nucleation and elongation occur. At small gating factors, both processes take place over similar time periods. Within this gating regime, small changes in the gating factor produce measurable changes in the number and lengths of filaments assembled over the course of the polymerization reaction. At large gating factors, the time periods for nucleation and elongation diverge. At the same time, the number and lengths of the polymerized actin filaments become much less sensitive to changes in the gating factor. These findings suggest that variation of the filament elongation rate produces the most significant changes to the distribution of actin monomers under conditions in which filament nucleation and elongation take place over similar periods of time.

Formin significantly increases the rate of filament nucleation in reactions containing profilin (Figure 8). The magnitude of this change in nucleation increases as the concentration of profilin increases. As a result, both the rate and the period over which nucleation occurs become less sensitive to profilin in the presence of formin. These effects on nucleation result in an increase in the number of filaments assembled over the course of each polymerization reaction, with the most dramatic changes occurring at profilin concentrations exceeding 5 µM. Formin-bound filaments elongate at rates that exhibit biphasic dependence on the concentration of profilin [17, 18]. Consistent with this dependence, filaments assembled in reactions containing formin and profilin attain equilibrium lengths that are approximately proportional to their initial elongation rates.

### A regime for the assembly of long formin-bound filaments

Our simulations and *in vitro* experiments demonstrate that reaction conditions dictate not only the rate at which polymerization proceeds, but also the final products of the reaction. By introducing formin into our polymerization reactions, we found that an increase in the nucleation rate produces more filaments at the expense of filament length. We also found that monomer sequestration is an effective way to increase filament length by favoring binding to a pre-existing barbed end over nucleation. As such, we propose that suppression of filament nucleation is a more efficient mechanism for the assembly of long filaments than an increase in the elongation rate. This strategy is consistent with our *in vitro* observation that Bni1p FH1FH2 assembles filaments that are only 20-50% longer than filaments polymerized by Bni1p FH2, despite mediating 4-times faster elongation (Figure 9C) [17, 34].

This mechanism also explains the assembly of very long formin-bound filaments (>20 µm) in elongation experiments monitored by TIRF microscopy [10, 17, 25]. These reactions utilize a low concentration of formin (typically <5 nM) and a concentration of actin monomers that is optimized to ensure a low rate of spontaneous nucleation. Elongation along a surface also protects filaments from shearing. In contrast to these traditional elongation studies, we did not see a measurable increase in the number of long filaments in our bulk *in vitro* reactions containing low concentrations of formin. However, it is likely that the relatively fast spontaneous nucleation rate at 2 µM actin, coupled with an increased probability of breakage for long filaments, shortened the average filament lengths in our assays.

In cells, actin filament lengths are regulated both by the availability of polymerization-competent monomers and by the specific elongation factors that are associated with each filament [5]. Unregulated formin-mediated polymerization would rapidly deplete the concentration of actin monomers and produce deleterious effects on cytoskeletal dynamics. However, autoinhibition keeps the concentration of active formin low [6] and saturation of actin monomers with profilin and other actin-sequestering proteins inhibits formin’s intrinsic nucleation efficiency [44]. In combination, these strategies enable the assembly of a small number of formin-bound filaments that can elongate rapidly and efficiently. Under these conditions, variations in the elongation rates mediated by different formin isoforms are most likely to impact the lengths of the filaments they assemble (Figure 10).

**Figure 10.**
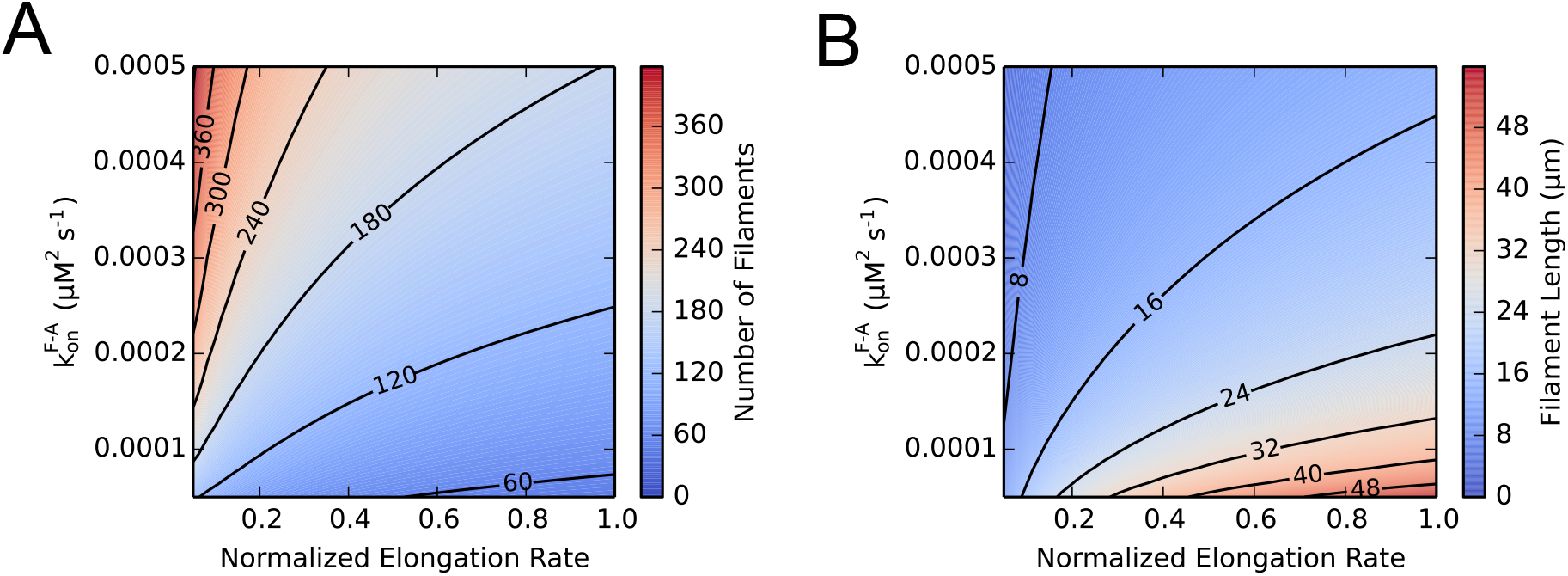
A low nucleation rate amplifies the influence of the elongation rate on filament length. Contour plots depicting the dependence of (A) the number and (B) the equilibrium lengths of filaments assembled in simulated polymerization reactions on the rate at which formins nucleate filaments via FH2-mediated actin monomer binding (*k*^F-A^_on_) and the elongation rate mediated by the formin. The elongation rate was varied by altering the gating factor and normalized to the maximal observed elongation rate. Simulations were carried out in the presence of 2 µM actin monomers, 1 nM formin and 5 µM profilin. Contour lines are shown at intervals of 60 filaments in (A) and 8 µm in (B).

### Use of kinetic modeling to measure affinities of formins for filament nuclei

Visualization of *in vitro* actin polymerization reactions captured the trends predicted by our simulations on a phenomenological level (Figures 3, 5 and 9). Introduction of formin into our bulk reactions produced a hyperbolic increase in the number of filaments and decrease in the average filament lengths (Figure 5B and C). Consistent with our simulations, the magnitude of these effects on filament number and length depended on the gating factor. The effects of Cdc12p on filament lengths closely matched our simulations performed at a gating factor of 0.05. However, Bni1p produced less dramatic changes in filament distribution than expected. This suggests that the rates we use to define the interactions between formin, actin and profilin more closely match the binding properties of Cdc12p than Bni1p. Whereas we used experimentally measured values for each formin’s elongation rates [17], the interactions between formins and actin nuclei during filament nucleation are not as well characterized. Thus, it is likely that Bni1p binds nuclei less tightly than we predicted. This discrepancy reveals a possible application for our model in fitting experimental data sets to extract the affinities of formin isoforms for filament nuclei. This could enable future dissection of the polymerization activities of formin isoforms that possess similar elongation activities but are known to stimulate nucleation at different rates [26, 27]. These formins would be predicted to generate actin filaments whose lengths and numbers depend both on their gating factor and on their intrinsic affinity for actin nuclei.

## Author Contributions

M.E.Z. designed research, performed research, analyzed data, and wrote the manuscript. L.A.S and B.M. performed research and analyzed data. N.C. designed research, analyzed data, wrote the manuscript, and acquired funding.

## Acknowledgements

This work was supported by National Institutes of Health (NIH) research grant R01 GM122787.

## Supporting Citations

References (46–50) appear in the Supporting Material.

