## Supplemental Materials for "Formin’s nucleation activity influences actin filament length"

| **Supplemental Table S1. Kinetic model parameter values** | | | | |
|---|---|---|---|---|
| **Reaction** | **Reaction Description** | | **Forward rate** | **Reverse rate** |
| Profilin-Actin Equilibrium | | | | |
| 1 | Profilin binds actin[a] | P + A = PA | 30 $\mu M^{-1}s^{-1}$ | 90 $s^{-1}$ |
| Free Filament Equilibria | | | | |
| *Nucleation of free actin monomers* | | | | |
| 2 | Monomer dimerization[b] | A + A = 2A | 35.7 $\mu M^{-1}s^{-1}$ | $1.63*10^{8}$ $s^{-1}$ |
| 3 | Monomer trimerization[b] | 2A + A = 3A | 2.18 $\mu M^{-1}s^{-1}$ | 1300 $s^{-1}$ |
| 4 | Free filament formation (addition of A)[b] | 3A + A = Free_BE | 9.6 $\mu M^{-1}s^{-1}$ | 0 |
| 5 | Free filament formation (Addition of PA)[c] | 3A + PA = Free_BE-P | 8 $\mu M^{-1}s^{-1}$ | 0 |
| *Free filament elongation* | | | | |
| 6 | Free actin incorporation at free barbed end[d] | Free_BE + A = Free_BE + FA | 11.6 $\mu M^{-1}s^{-1}$ | 1.4 $s^{-1}$ |
| 7 | Profilin-actin incorporation at free barbed end[e] | Free_BE + PA = Free_BE-P + FA | 10 $\mu M^{-1}s^{-1}$ | Detailed balance |
| 8 | Free actin incorporation at free pointed end[d] | Free_PE + A = Free_PE + FA | 1.3 $\mu M^{-1}s^{-1}$ | 0.8 $s^{-1}$ |
| *Profilin binds to free barbed ends* | | | | |
| 9 | Profilin binds to free barbed end[f] | Free_BE + P = Free_BE-P | 10 $\mu M^{-1}s^{-1}$ | 2500 $s^{-1}$ |
| Formin-Bound Filament Equilibria | | | | |
| *Formin-mediated nucleation* | | | | |
| 10 | Formin-mediated nucleation[g] | FH1-FH2 + A + A = Bound_PE | 0.0002 $\mu M^{-1}s^{-1}$ | 0 |
| 11 | Formin-mediated nucleation (A + PA)[g] | FH1-FH2 + A + PA = Bound_BE-P | 0.0002 $\mu M^{-1}s^{-1}$ | 0 |
| *Profilin binds to formin-bound barbed ends* | | | | |
| 12 | Profilin binds to bound barbed end[f] | Bound_BE + P = Bound_BE-P | 10 $\mu M^{-1}s^{-1}$ | 2500 $s^{-1}$ |
| *FH2-mediated elongation of formin-bound filaments* | | | | |
| 13 | Actin incorporation at bound barbed end[h] | Bound_BE + A = Bound_BE + FA | p*11.6 $\mu M^{-1}s^{-1}$ | p*1.4 $s^{-1}$ |

| | | | | |
|---|---|---|---|---|
| 14 | Profilin-actin incorporation at bound barbed end[i] | Bound_BE + PA = Bound_BE-P + FA | p*10 $\mu M^{-1}s^{-1}$ | Detailed balance |
| *FH1-mediated elongation of formin-bound filaments* | | | | |
| 15 | Profilin binds to FH1[j] | FH1 + P = FH1-P | 200 $\mu M^{-1}s^{-1}$ | 5500 $s^{-1}$ |
| 16 | Profilin-actin binds to FH1[k] | FH1 + PA = FH1-PA | 40 $\mu M^{-1}s^{-1}$ | Detailed balance |
| 17 | Actin binds to profilin-bound FH1[l] | FH1-P + A = FH1-PA | 20 $\mu M^{-1}s^{-1}$ | 60 $s^{-1}$ |
| 18 | FH1 loop closure[m] | FH1_o = FH1_c | $r_0 * n_r^{-3/2}$ $s^{-1}$ | Detailed balance |
| 19 | FH1-profilin loop closure[m] | FH1-P_o = FH1-P_c | $r_0 * n_r^{-3/2}$ $s^{-1}$ | 2500 $s^{-1}$ |
| 20 | FH1-profilin-actin loop closure[n] | FH1-PA_o = FH1-P_c + FA | p* $r_0 * n_r^{-3/2}$ $s^{-1}$ | Detailed balance |
| *Pointed end elongation of formin-bound filaments* | | | | |
| 21 | Free actin incorporation at pointed end of formin-bound filament[d] | Bound_PE + A = Bound_PE + FA | 1.3 $\mu M^{-1}s^{-1}$ | 0.8 $s^{-1}$ |
| General Reaction Parameters | | | | |
| | Profilin-actin dissociation constant[o] | $K_d$_PA | 3 µM | |
| | FH2 gating factor[p] | p | 0.5 | |
| | Kinetic prefactor[q] | $r_0$ | $10^5$ | |
| | Distance (in residues) from profilin binding site in FH1 to barbed end | $n_r$ | 22 | |

[a]The association rate was used in simulations performed by Paul and Pollard (1) and estimated from Vinson and coworkers (2). The dissociation rate was calculated based on the dissociation constant of 3 µM for actin and *S. cerevisiae* profilin (3).

[b]Rates from Sept and McCammon (4). A filament is established via the association of an actin monomer with an actin trimer, and we treat this reaction as irreversible, consistent with the studies of Sept and McCammon and Paul and Pollard (1, 5).

[c]The association rate was used in the simulations performed by Vavylonis and coworkers and Paul and Pollard (1, 6). We treat this reaction as irreversible, consistent with the studies of Sept and McCammon and Paul and Pollard (1, 5).

[d]Rates measured by Pollard (7).

[e]These rates were used by Vavylonis and coworkers (6). The reverse rate was obtained from detailed balance calculations.

[f]These rates were used by Vavylonis and coworkers and Paul and Pollard (1, 6). The dissociation rate was derived using the affinity of human profilin 1 for filament barbed ends (~250 µM (8, 9)). It also assumes that the FH2 domain does not influence the affinity of profilin for the barbed end (6).

[g]As in the Paul study, we assume that formin irreversibly nucleates a filament by binding either two actin monomers or one actin monomer and one profilin-actin complex (1). Although formin likely binds the actin monomers and/or profilin-actin sequentially, we simplified this process to a trimolecular reaction.

[h]Association and dissociation rates derived from Pollard (7) and multiplied by the FH2 gating factor "p"(6).

[i]These rates were used by Vavylonis and coworkers.

[j]These rates were used by Paul and Pollard. The association rate is based on an estimation that the association rate constant of bovine spleen profilin with poly-L-proline oligomers exceeds 200 $\mu M^{-1}s^{-1}$ (10) and an assumption that polyproline tracts in FH1 domains are shorter and less mobile than the oligomers used by the Perelroizen study (6). The dissociation rate was calculated from the equilibrium dissociation constant of *S. cerevisiae* profilin from polyproline (27.5 μM) (3). For simplicity, our model includes only one polyproline tract in the formin's FH1 domain.

[k]The association rate is slower than the rate of profilin:FH1 association to account for the decreased mobility of profilin-actin compared to free profilin (6).

[l]The association rate was used by Vavylonis and coworkers (6). It was selected based on published rates of 35-53 $\mu M^{-1}s^{-1}$ for the associations of amoeba profilin II and bovine spleen profilin with actin (2, 10), and accounts for the reduced mobility of FH1-bound profilin. Perelroizen and coworkers also demonstrated that free profilin and polyproline-bound profilin bind actin monomers with the same affinity (10). Likewise, our calculation of the dissociation rate assumes that tethering profilin to the FH1 domain does not influence its affinity for actin.

[m]This rate was used by Vavylonis and coworkers and assumes the FH1 domain is a random coil and that the binding of FH1-bound profilin to the barbed end is not influenced by FH2 gating (6).

[n]This rate was used by Vavylonis and coworkers and accounts for the effects of FH2 gating on the binding of actin monomers to formin-bound barbed ends (6).

[o]This dissociation constant was measured by Eads and coworkers (3).

[p] This value was first reported by Kovar and coworkers (11).

[q]This parameter is a microscopic rate into which all orientational effects have been collapsed (6). The rate of association of the two ends of an unfolded peptide is $10^7$ $s^{-1}$ (12). Our rate is slower to account for the probability of collisions between FH1-tethered profilin and the barbed end occurring in the correct orientation to promote a binding event.

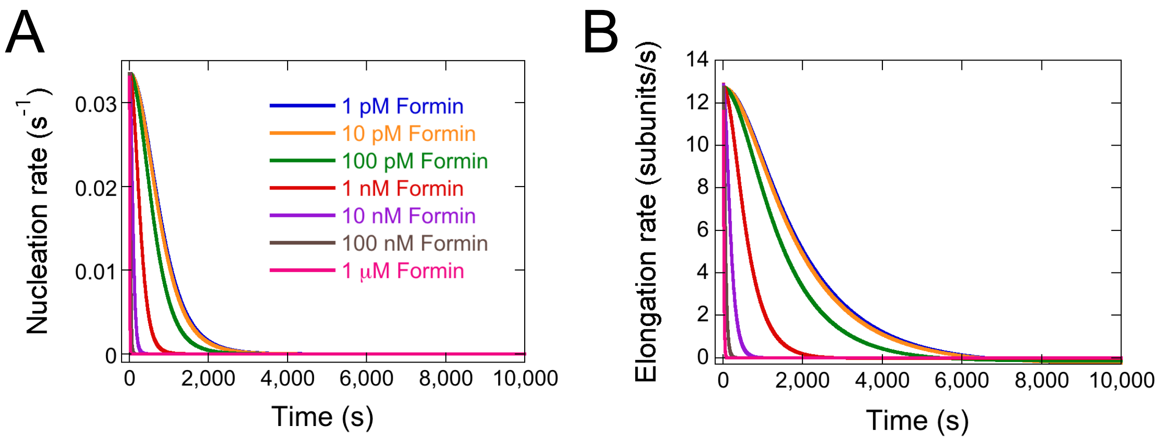


**Supplemental Figure S1. Assembly of filaments with free barbed ends in the presence of formins.**

Quantification of simulated polymerization reactions containing 2 µM actin, 1 nM formin and 0 µM (blue lines), 0.5 µM (orange lines), 1 µM (green lines), 2.5 µM (red lines), 5 µM (purple lines), 10 µM (brown lines) or 25 µM (pink lines) profilin. Profilin's affinity for actin monomers was set at 3 µM. Each simulation was carried out over 10,000 s. (A) Spontaneous filament nucleation rate over time. (B) Rate of elongation of filaments with free barbed ends over time.
